# Environmental Niche Adaptation Revealed Through Fine Scale Phenological Niche Modeling

**DOI:** 10.1101/534636

**Authors:** Matthew H. Van Dam, Andrew J. Rominger, Michael S. Brewer

## Abstract

**Aim:** Phenology, the temporal response of a population to its climate, is a crucial behavioral trait shared across life on earth. How species adapt their phenologies to climate change is poorly understood but critical in understanding how species will respond to future change. We use a group of flies (*Rhaphiomidas*) endemic to the North American deserts to understand how species adapt to changing climatic conditions. Here we explore a novel approach for taxa with constrained phenologies aimed to accurately model their environmental niche and relate this to phenological and morphological adaptations in a phylogenetic context.

**Taxon:** Insecta, Diptera, Mydidae, *Rhaphiomidas*

**Location:** North America, Mojave, Sonoran and Chihuahuan Deserts.

**Methods:** We gathered geographical and phenological occurrence data for the entire genus *Rhaphiomidas*, and, estimated a time calibrated phylogeny. We compared Daymet derived temperature values for a species adult occurrence period (phenology) with those derived from WorldClim data that is partitioned by month or quarter to examine what effect using more precise data has on capturing a species’ environmental niche. We then examined to what extent phylogenetic signal in phenological traits, climate tolerance and morphology can inform us about how species adapt to different environmental regimes.

**Results:** We found that the Bioclim temperature data, which are averages across monthly intervals, poorly represent the climate windows to which adult flies are actually adapted. Using temporally-relevant climate data, we show that many species use a combination of morphological and phenological changes to adapt to different climate regimes. There are also instances where species changed only phenology to track a climate type or only morphology to adapt to different environments.

**Main Conclusions:** Without using a fine-scale phenological data approach, identifying environmental adaptations could be misleading because the data do not represent the conditions the animals are actually experiencing. We find that fine-scale phenological niche models are needed when assessing taxa that have a discrete phenological window that is key to their survival, accurately linking environment to morphology and phenology. Using this approach, we show that *Rhaphiomidas* use a combination of niche tracking and adaptation to persist in new niches. Modeling the effect of phenology on such species’ niches will be critical for better predictions of how these species might respond to future climate change.

## INTRODUCTION

Evaluating species response to climate change and predicting future distributions is of concern to ecologists, evolutionary biologists and policy makers interested in preserving biodiversity (Parmesan, 2006). In addition, identifying whether a particular species or population adapts to climate change by adjusting its phenology or morphological/physiological traits, as opposed to changing its range via dispersal, is key to predicting future distributions, extinctions (Holt, 1990) and community disassembly (Sheldon et al., 2011). Dispersal is often assumed to be the primary response of populations and species to climate change (Thomas et al., 2004); however, this might not be the case (Visser, 2008). Dispersal limitation and heterogeneous or fragmented landscapes could preclude spatial tracking of climatic niches (Hof et al., 2011). In such cases, populations must adapt their phenology and/or morphology in order to persist (Hoffmann & Sgrò, 2011). Such adaptation, however, is often not accounted for in models of species’ responses to climate change (Bradshaw & Holzapfel 2008; van Asch et al. 2007).

Phenology plays a critical role in the reproductive cycles and developmental processes of many organisms (Mitchell et al., 2008; Banta et al., 2012; Zohner & Renner, 2014; Gerst et al., 2017; Zhang & Hepner, 2017). The past century of climate change has already produced measurable changes in phenology, especially in plants (Ellwood et al., 2013; Everill et al., 2014; Zohner & Renner, 2014). Therefore, finding a better way to model critical environmental conditions linked to phenological cycles and examining how different species adapt their phenologies is important for understanding species’ ecologies and their conservation. Phenological responses have been identified and tested through long-term ecological studies (Miller-Rushing & Primack, 2008) but can also be understood in a phylogenetic context along with morphological evolution.

Here, we use a group of flies (Diptera: Mydidae: *Rhaphiomidas* Osten Sacken, 1877; Fig. 1) endemic to the North American deserts to understand how species adapt to changing climatic conditions. Specifically, we are interested in whether the flies change their emergence time to track the same basic seasonal climate window, or whether the adult flies adapt physiologically/morphologically to changing climate during that window.

**Figure 1.**
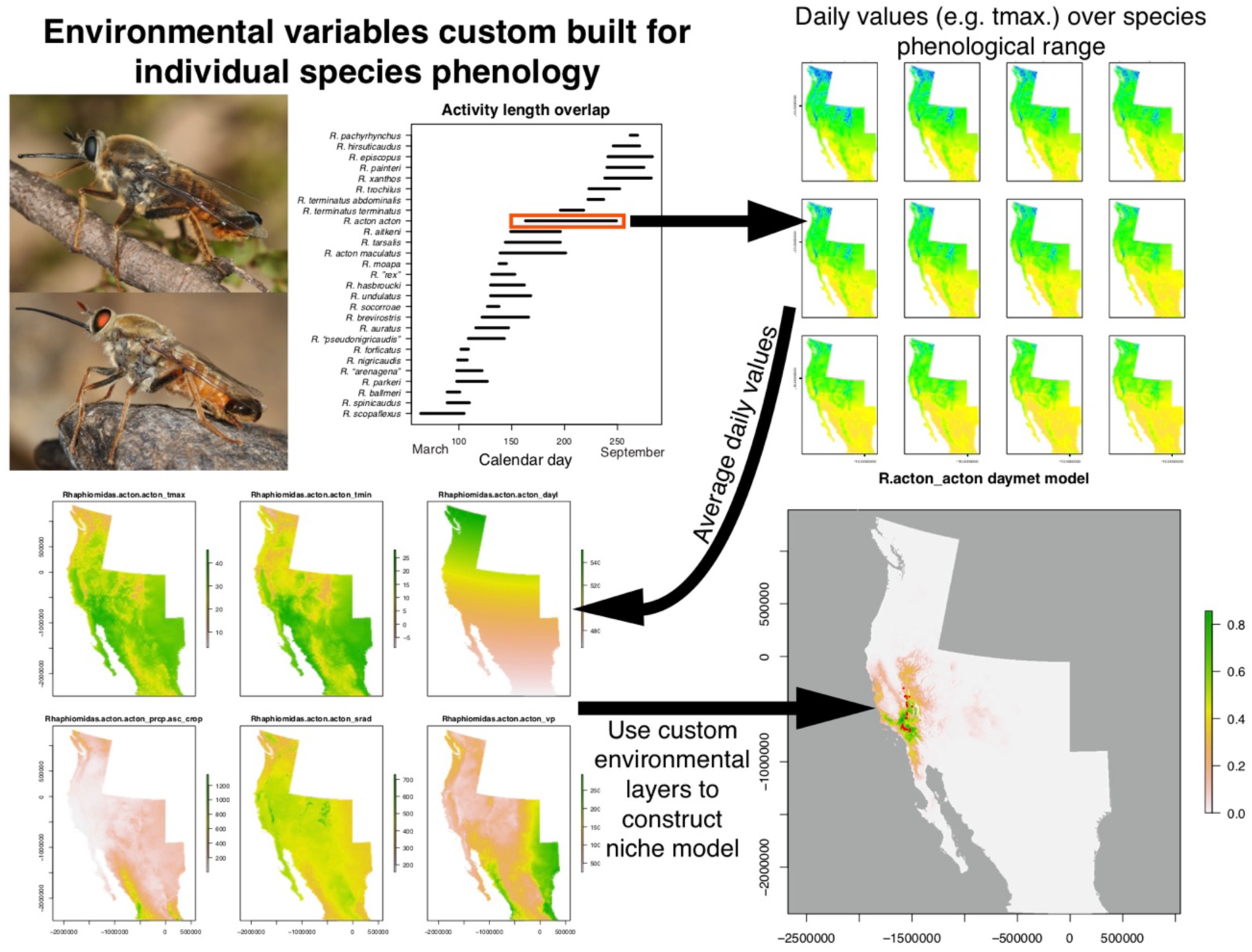
**Upper left**, *Rhaphiomidas pachyrhynchus* adult male, *R. sp. “pseudonigricaudis”* adult male. **Central panel**, Activity duration of *Rhaphiomidas* species. **Upper right**, daily values over a species’ adult activity length. **Lower left**, resulting raster layers for each environmental variable averaged or summed in the case of precipitation over the activity length period of a species. **Lower right**, resulting environmental niche model from custom raster layers for a species.

The genus *Rhaphiomidas* is distributed throughout the deserts of the Southwestern United States and Northern Mexico (Cazier, 1985; Van Dam, 2010). Most species of *Rhaphiomidas* feed on floral nectar as adults and thus phenological matching with their plant resources may be critical for both mutualists. Species are either restricted to aeolian (wind-blown sediments) sand dunes (15/27 species) or loose alluvial sands (Van Dam, 2010), with many species (9/27) endemic to a single dune system. *Rhaphiomidas* adults fly in the spring and fall and are most active during spring and fall blooms. Some species do not appear to feed as adults, making the synchronization of their activity all the more important for reproductive success. Multiple studies have been conducted on the behavior of adult flies, noting the temperatures requirements for adult flight activity (Toft & Kimsey 1982; Ballmer *et al.* 1994; Rogers & Mattoni 1993; Kingsley 1996). Larvae and pupae of *Rhaphiomidas* are entirely subterranean at depths of up to 1.5 m (M.H.V.D pers. obs.) and thus only experience the ambient climate for the short duration of their lives spent above ground. This dramatically discrete phenology makes them ideal for studying phenological response to climate.

Studies of climatic niches typically depend on species distribution models to understand the correlation between a species’ occurrences and underlying climatic variables (Warren et al., 2018). Such studies rarely incorporate phenology when reconstructing niche occupancy, often using only monthly averaged WorldClim data (Hijmans et al., 2005). Constraining climate data to monthly averages may misrepresent the climate that species are actually experiencing. Although the WorldClim variables are appropriate for capturing realized niches based on abiotic environmental factors experienced year-round, such as for perennial plants and some mammals, reliance on these variables could lead to biased predictions for an entire suite of organisms that have discrete phenologies.

To solve this problem, we used climate data that were averaged by day at a 1km square scale available from Daymet (Thornton et al. 2014). To measure how species adapt to different climate conditions, the Daymet-based approach was put in a phylogenetic framework to test whether phenological adaptation (niche tracking) or physiological/morphological adaptation (niche adapting) has occurred. A phylogenetic framework allows us to account for the role of shared evolutionary history (Felsenstein 1985) in producing the observed correlations between traits and niche preference. We can also infer the evolutionary history of adult niche preference and measure the rate at which changes have occurred.

Understanding how species adapt to climate change in different climate regimes by adapting their morphology and physiology is also important for mitigating extinction risk due to climate change (Visser, 2008). For many species that rely on ambient temperature for thermoregulation, climate change is predicted to have a negative effect on their survival (Quintero & Wiens, 2013). In general, species or populations that are poikilotherms (commonly known as cold-blooded) have a darker coloration in colder environments (Shapiro, 1976; Kingsolver & Wiernasz 1991). For example, reptiles become darker in colder conditions to maintain activity (Bittner et al., 2002; Rosenblum, 2005). This adaptation helps them achieve metabolic activity more quickly. We propose that the dark coloration seen in some *Rhaphiomidas* species is an adaptation facilitating absorption of more solar energy, thereby allowing increased activity in cooler climates.

## METHODS

### Analysis Pipeline

In order to test for phenological changes that track a niche and/or if species adapt to environmental conditions, we analyzed a diverse set of data (environmental and phylogenetic). The methods can be broken down into five parts:

1. We processed environmental variables and constructed niche models.
2. We created predicted niche occupancy profiles (PNO, in particular thermal maximum) to relate the physical temperature corresponding to probability densities of the niche model (Elith et al., 2011).
3. We constructed a time-calibrated phylogeny for *Rhaphiomidas*.
4. Using the phylogeny, we tested whether species possessing dark coloration are correlated with colder thermal maxima, indicating an adaptive response to colder temperatures and examined ancestral state reconstruction using phylogenetic comparative methods to ensure this wasn’t an artifact of inheritance.
5. We combined phylogenetic reconstructions of traits (morphology, phenology and temperature) to see if sister species changed their phenology relative to changes in temperature and morphology.

### DNA Data and Alignment

*Rhaphiomidas* sequences were taken from Van Dam & Matzke (2016). Contigs were assembled and edited in Geneious Pro v. 4.6.4 (Biomatters Ltd.). Sequences were aligned by ClustalW-2.0.10 (Larkin et al., 2007) with parameters set to GAPOPEN=90.0, GAPEXT=10. Sequences were colored by amino acid in Mesquite version 2.71 (Build 514) (Maddison & Maddison 2009) to check for stop codons.

### Phylogenetic Analyses

We obtained data for 219 individuals, including 183 *Rhaphiomidas* exemplars and 36 outgroup samples. These data comprised 2904 bp of mtDNA (COI, COII, and 16S genes), and 3720 bp of nDNA (EF1alpha, PGD, snf, Wg, and CAD). The best partitioning scheme was determined in PartitionFinder 1.1.1 (Lanfear et al., 2012) using the ‘greedy’ algorithm. The partitioning scheme was selected using the Akaike information criterion corrected for sample size (AICc). Relationships among *Rhaphiomidas* species were reconstructed using Bayesian Inference (BI) implemented in MrBayes 3.2 (Ronquist et al., 2012). Reversible-jump MCMC was used to explore the entire model space of the general time reversible (GTR) substitution models for our different character sets (Huelsenbeck et al., 2004). We ran two independent runs comprising eight total Markov Chains for 70 million generations, sampling every 1000th generation. Split-frequencies and log-likelihood curves were examined in Tracer 1.5.3 (Rambaut & Drummond 2018).

The first 35,000 of the 70,000 trees sampled were removed as burnin. Standard deviation of split-frequencies <0.05 between the two MrBayes runs suggested reasonable convergence. A maximum clade credibility tree was constructed with *DendroPy SumTrees* function (Sukumaran & Holder, 2010) using median edge lengths. This tree was inspected for evidence of mitochondrial introgression as in McGuire et al., (2007). The dataset was then pruned to a single specimen per species, preferring specimens with the most complete sequence that did not show signs of mitochondrial introgression. This was done because Yule and Birth Death (BD) tree priors in BEAST treat every tip as a species; a tree with multiple specimens for each species can result in a biased date inference (Drummond & Bouckaert, 2015). The MrBayes tree was pruned to the same specimens, and the pruned dataset was analyzed in BEAST version 1.8.2 (Drummond et al., 2012) with the pruned MrBayes tree serving as the starting topology for phylogenetic dating analyses. We used stepping stone (SS) sampling (Drummond et al., 2016) to select the tree prior (Birth-Death chosen over Yule) and clock model (a lognormal relaxed clock chosen over strict clock). The MCMC process was run for 4×10^7^ generations sampling every 1000 generations. Stationarity was assessed using Tracer version 1.5.3. The effective sample size (ESS) for model parameters was also examined to assess the adequacy of the post-burnin samples. The tree was calibrated using a fossil calibration point for the origin of the Mydinae in the late Cretaceous (Martill et al., 2007) using a normal distribution with a mean of 120 Ma and a standard deviation of ±10 Ma.

### Data Acquisition/Niche Modeling

Specimen data for *Rhaphiomidas* were acquired from museum collections (EMEC, CAS, UCR, LACM) and previously published literature (Cazier 1985; Rogers and Mattoni 1993; Rogers and Van Dam 2007; Van Dam 2010), as well as from the personal collection of M.H.V.D. Locations were georeferenced in ArcGIS (ESRI 2011) and were recorded in both WGS_84 (degrees-minutes-seconds) and Lambert Conformal Conic projection required for Daymet georeferencing (http://daymet.ornl.gov/datasupport). A total of 409 occurrence points were included, with an average of 15 per species, median 13 and a range of 10–40. Several species are endemic to a single dune system, so 10 points were randomly scattered across the spatial extent of such dune fields. This number was selected because it has been demonstrated to be a reasonable minimum number of points needed for species distribution models using Maxent (Hernandez et al., 2006; Evans et al., 2009, van Proosdij et al. 2016). To obtain the most accurate possible temperature estimates when adult flies are active, we extracted climate data from the Daymet climate data base (Thornton et al. 2014). This allowed us to partition climate data by day and average over a species’ occurrence time (typically weeks) rather than using monthly averages. A list of the Daymet tiles is provided in the Supporting Information. The 1 km by 1 km resolution tiles were downloaded from years 1980–2011. A total of six variables were used from the Daymet database (vapor pressure *vp*, day length *dayl*, precipitation *prcp*, solar radiation *srad*, temperature max *tmax* and temperature minimum *tmin*). The data totaled just over 2 terabytes of information. The first and last occurrence times were recorded from the specimen data (Fig. 1) and used to define the time slices to extract from the Daymet climate data.

Extracted environmental variables were averaged over the 30 years. Precipitation data were summed by time slice and then averaged across years. This procedure was performed in R statistical software with a custom script (see Supporting Information). We utilized the R packages *ncdf4*, *raster*, *maps* and *dismo* for this process (Hijmans & van Etten, 2012; Becker et al., 2017; Hijmans et al., 2017; Pierce, 2017). Niche models were constructed in MaxEnt (Phillips et al., 2006) using the R package *dismo*. We examined collinearity of predictors using *SDMTools* (Warren et al., 2018; Warren et al., 2010).

To evaluate the extent to which Daymet and BioClim representations of the same geographic point diverge, we extracted temperature values from each respective raster layer for the same point. Because *Rhaphiomidas* are late spring and summer active species, we compared the values of the tmax (Daymet) data with Bioclim BIO5 (Max Temperature of Warmest Month) and BIO10 (Mean Temperature of Warmest Quarter).

### Predicted Niche Occupancy

To identify how *Rhaphiomidas* lineages adapted to change in different climate regimes, we constructed each species' predicted niche occupancy (PNO) profile from the cumulative probability of occurrence against the environmental data (Evans et al., 2009). We then used the PNO to calculate the weighted mean for each species. PNO profiles were constructed in *phyloclim* (Heibl & Calenge, 2013), and 100 random samples were drawn from the PNO profile as in Evans et al. (2009) to calculate the weighted mean. Calculations for PNOs from the Daymet data were derived using a slightly different method than those for the Bioclim data. Because each one of our Daymet environmental layers is unique to each taxon, some values were not found among layers. Thus, we first calculated the range across all individual taxa layers for a variable and then merged the PNO files by taking the floor to the nearest °C. For instance, 32.5456845°C in species-A PNO and 32.59975962°C in species-B PNO were changed to 32°C in the two PNOs, so they can then be used for calculating the weighted means as above.

### Evaluating the correlation between temperature and body color of Rhaphiomidas

We categorized fly coloration according to the following rules:1) if the first 3 abdominal tergites were almost entirely black, they were coded as dark, 2) if the first 3 abdominal tergites were orange or silver they were coded as pale. Because there were no intermediate states, we treated these as discrete.

To examine whether colder daytime temperatures are correlated with a darker body color while accounting for shared evolutionary history, we used a threshold model from quantitative genetics (Wright, 1934; Felsenstein, 2012), using MCMC to sample the unknown liabilities in a postulated continuous trait underlying a discrete character. We used the R package *phytools* 0.3-72 (Revell, 2012), which implements this model in the function *threshBayes*. To account for differences in branch lengths and topologies from our posterior distribution of trees, we sampled 10% of the posterior trees resulting in 3900 sampled trees. We then measured the correlation between the two characters over this set of trees. We ran the MCMC chain for 5×10^6^ generations sampling every 1,000 generations for each tree. After examining the trace plot of the posterior, we set a burn-in of 10%.

### Identifying Coloration Shifts

To identify shifts in coloration, we reconstructed ancestral states for coloration on the phylogeny. First, we sorted the trees from the BEAST posterior (minus the burnin) according to their topologies (grouping identical topologies together). We used the *DendroPy SumTrees* function (Sukumaran & Holder, 2010), taking median edge lengths, to create a representative strict consensus tree for each set of unique topologies. We then used a Bayesian threshold model implemented in *phytools* with the function *ancThresh*, running 2.5×10^5^ MCMC generations sampling every 1,000 with a burnin of 50 over each unique topology.

### Ancestral State Reconstructions and Evaluating Climate Disparity Through Time

To reconstruct ancestral states for the continuous traits of temperature and occurrence time (phenology), we first determined the best evolutionary model, Brownian Motion (BM) or Ornstein−Uhlenbeck (OU), given our tree and data using the *fitContinuous* function in *geiger* (Harmon et al., 2008). In instances where the OU model was selected, we used a modified version of the *phytools* function *contMap* where the internal *anc.ML* function was set to the OU model.

Ancestral state estimates were used to measure the relative amount of disparity between sister nodes using *geiger*'s disparity function to calculate the average squared Euclidean distance in each clade. To visualize the amount of change between nodes on the tree, we plotted a phenogram for both traits using the *phenogram* function of *phytools*. We also calculated the difference between parent and daughter node values (normalized temperature and phenology) from our *contMap* reconstructions using a custom R script utilizing functions from BioGeoBEARS (Matzke 2014). These values were used to color the branches and help visualize the change in trait values along the tree.

## RESULTS

### Molecular Data and Phylogenetics

The BEAST analysis indicated that *Rhaphiomidas* is sister to the remaining Mydidae, and these groups together are sister to the Apioceridae (see Supporting Information). All model parameters had an ESS (effective sample size) >200, suggesting sufficient sampling of the posterior distribution. The root node of *Rhaphiomidas* is estimated to have diverged 70 ± 42Ma (node height 95% HPD), with most of the species diversifying in the last 18.5 ± 11Ma. Terminals showing evidence of introgression were removed, leaving a topology that was also congruent with species concepts. From the Stepping Stone sampling results, we rejected the strict clock prior and Yule tree prior in favor of the relaxed lognormal clock prior and the Birth-Death tree model (Supporting Information).

### Niche Modeling Results

First, we compared the distributions of point values for temperature extracted from raster layers between the Daymet tmax data and Bioclim BIO5 and BIO10. We found that the Bioclim data did not adequately reflect the conditions that these species were experiencing as adults (Fig. 2). Many of the Bioclim values were outside of the species’ temperature range and further did not show differentiation between allochronic species, especially in the case of BIO5. Because the measurements of these two data sets are otherwise congruent (Hijmans et al. 2005), this difference is mostly due to the time periods binned in making the rasters. As the maximum temperature is key to understanding further hypotheses of morphological adaptations, we proceeded using the Daymet raster layers because they are more representative of the environmental conditions that these species experience.

**Figure 2.**
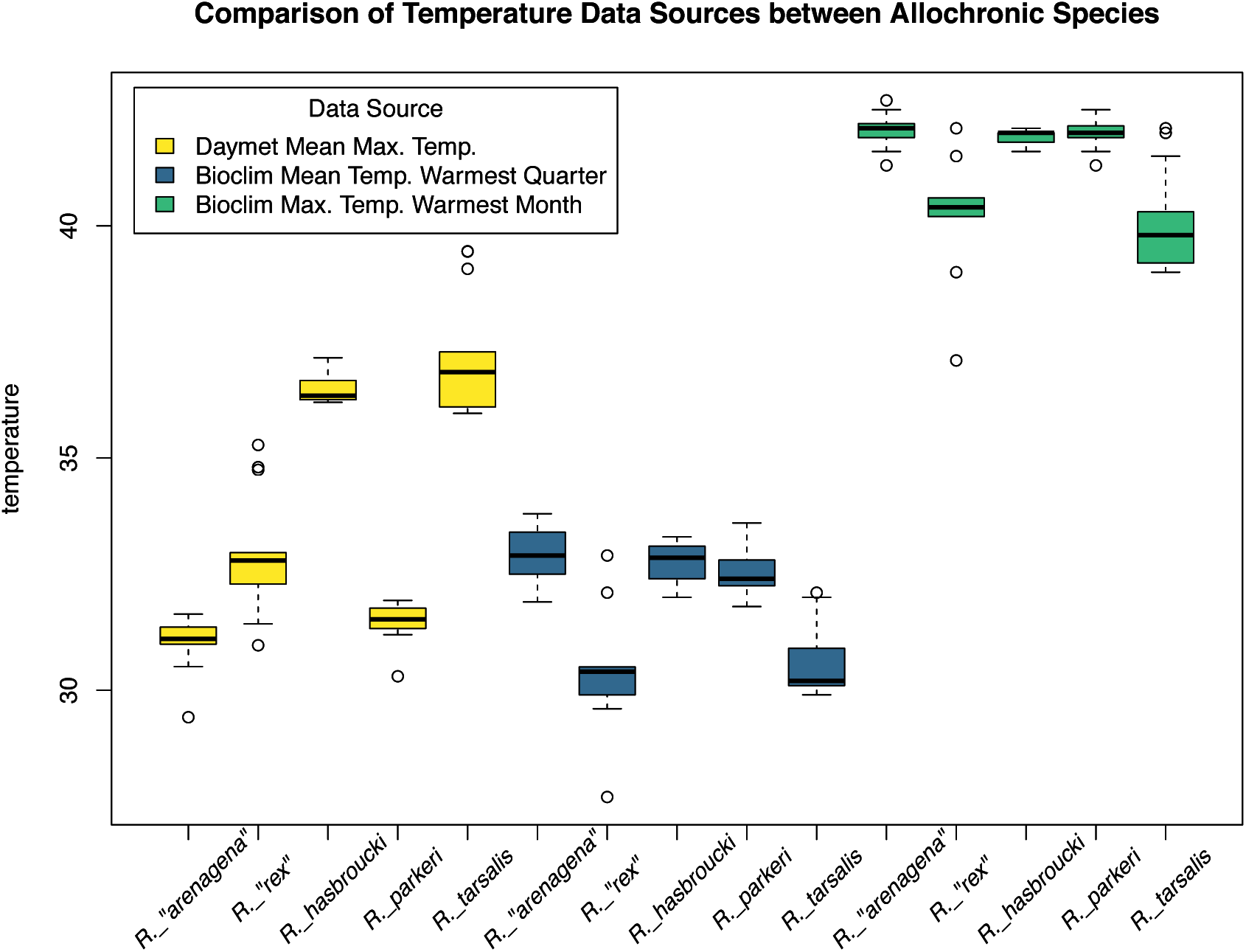
Box plots of temperature values extracted from occurrence points: Mean daily maximum temperature (tmax) from Daymet data (yellow), Bioclim Bio10 mean temperature warmest quarter (blue), Bioclim Bio5 maximum temperature warmest month (green). *R. “arenagena”* and *R. “rex”* are partially sympatric but emerge at different times (allochronic). *R. hasbroucki* and *R. parkeri* are also partially sympatric but are allochronic. *R. tarsalis* is the sister taxon to *R. hasbroucki*, they are allopatric and allochronic.

The collinearity between the predictor variables varied between strongly correlated (tmax and tmin) to highly uncorrelated (vapor pressure and solar radiation) (see Supporting Information). Given the small number of predictors and the biological importance of tmax and tmin we left both of these predictor variables in further analyses of species’ niches. The contribution of the six bioclimatic variables varied considerably between species. Variables, such as precipitation, consistently contributed to large percentages of the model probability – 90% in the case of *R. undulatus*. Other variables such as DayLength contributed modestly to the model predictions, ranging up to 30% for *R. xanthos*. AUCs varied from 0.65 to 1 (see Supporting Information). The weighted mean for tmax ranged from 25.1 – 38.5° C across species.

### Evaluating the Correlation between Temperature and Body Color

The mean correlation coefficient between color and temperature is −0.70 over all trees, indicating that as temperature decreases, there is a trait change from pale to dark (pale coded as 0 and dark coded as 1; Fig. 3). The 95% HPD varied from −0.93 to −0.25, and none of the individual trees examined had a 95% HPD that overlapped with zero (Supporting Information). Because this is a Bayesian analysis, significance is assessed through the 95% credible interval and its overlap with 0. Our 95% credible interval does not overlap with zero (Fig. 3).

**Figure 3.**
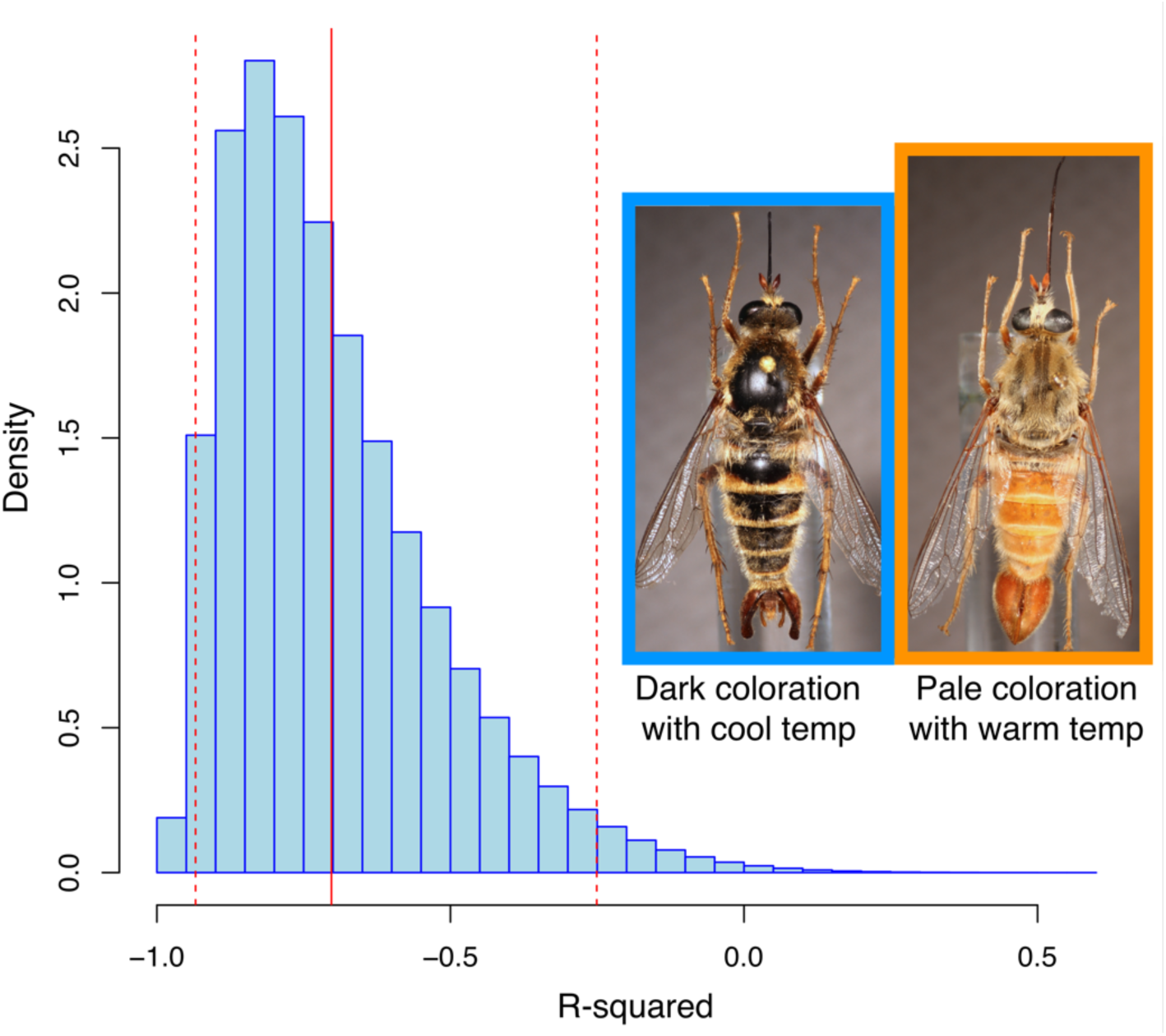
*threshBayes* posterior density distribution of the correlation between *Rhaphiomidas* body color (dark or light) and Daymet mean daily maximum temperature. The histogram shows the posterior distributions of 3900 trees (10% of the sampled trees), the solid red line indicates the grand mean, the dashed lines indicate the mean of the 95% confidence intervals. None of the individual 95% confidence intervals from the set of trees overlap with zero, indicating a significant correlation, robust to topology and branch lengths.

### Identifying Coloration Shifts

The common ancestor of *Rhaphiomidas pachyrhynchus* and *R. episcopus* was estimated to have dark coloration in all tree topologies. Only 2/48 topologies (representing only 3/3900 trees) had any other nodes reconstructed as dark in coloration. Thus, the ancestral states are likely pale for all but one node in the tree. The results of the maximum clade credibility tree are shown in Fig. 4.

**Figure 4.**
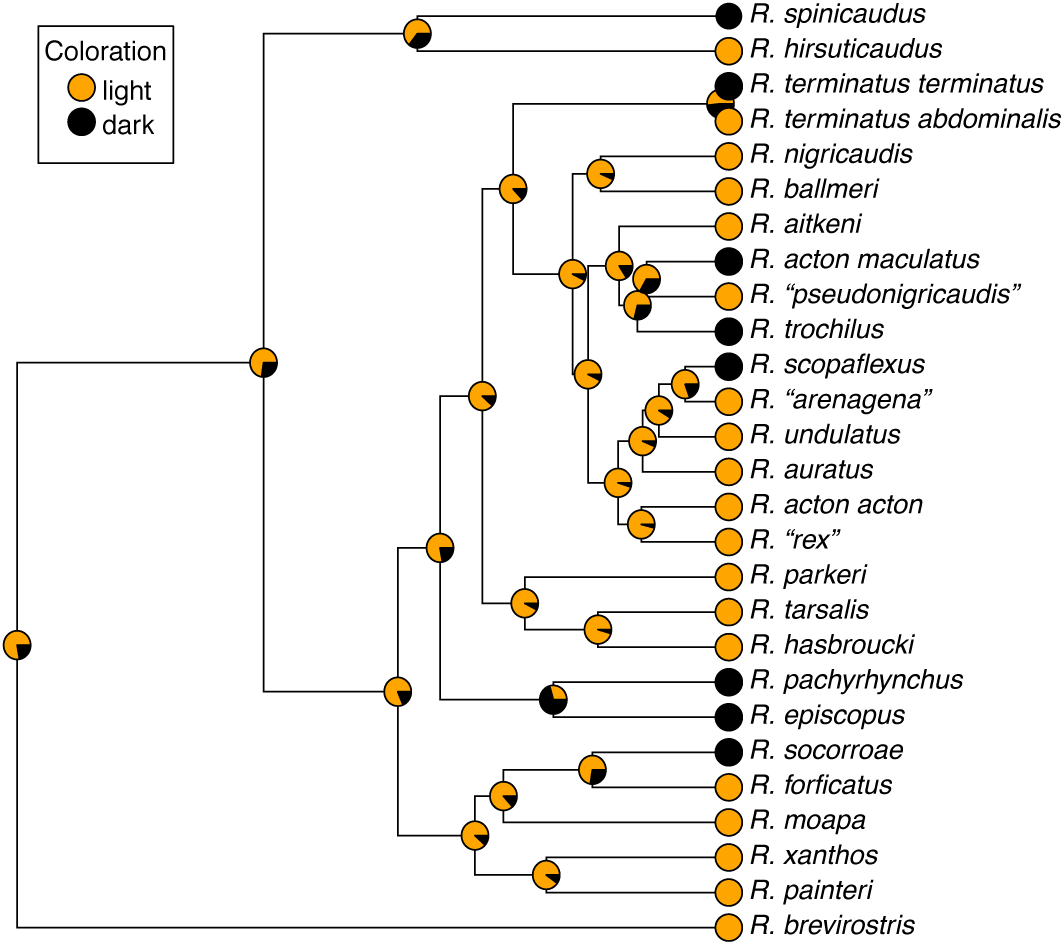
*Rhaphiomidas* maximum clade credibility tree, with ancestral state reconstructions for body coloration. Ancestral light or dark coloration was reconstructed via the *ancThresh* function in the *phytools* R package (Revell 2012). The analysis was run for 2.5×10^6^ MCMC generations, sampling every 1000, with a burnin of 50.

### Ancestral State Reconstructions and Evaluating Climate Disparity Through Time

The OU model was an overwhelmingly better fit than BM for weighted mean tmax dAICc [OU:0, BM:32.6278], but BM was slightly preferred for phenology dAICc [OU:0.441541, BM:0].

The results of the *contMap* reconstructions suggest most species of *Rhaphiomidas* tend to occupy a relatively warm climate space, and most species occur in late spring with only a few phenological shifts from spring to fall (Fig. 5). We find relatively few nodes where there is more disparity in temperature than in phenology (6 of 26), with 4 of 26 nodes showing approximately equal disparity between temperature and phenology and 16 of 26 nodes showing more disparity in phenology than in temperature (Fig. 6). In addition, the largest shifts in temperature occur at the terminal nodes in the tree, whereas large shifts in phenology occur at both internal and terminal nodes (Fig. 7).

**Figure 5.**
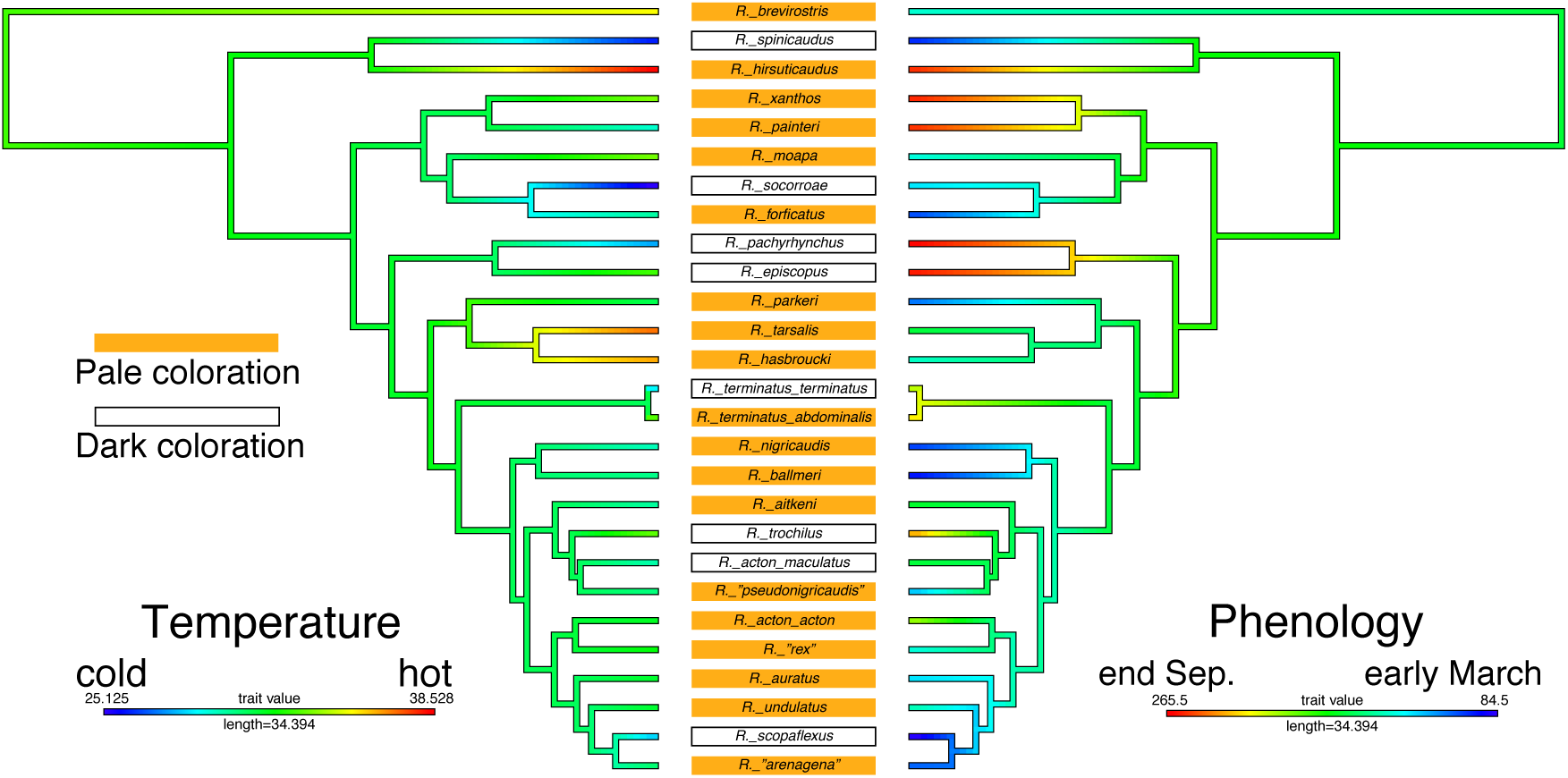
Results of the *contMap* ancestral state reconstructions. Left tree, ancestral state reconstructions for mean daily temperature maximum value, warmer colors along branches indicate warmer maximum temperatures. Right tree, ancestral state reconstructions for mid-occurrence times, warmer colors along branches indicate occurrence times later in the year. Orange rectangles indicate species with a light coloration, black boxes indicate species with a dark coloration.

**Figure 6.**
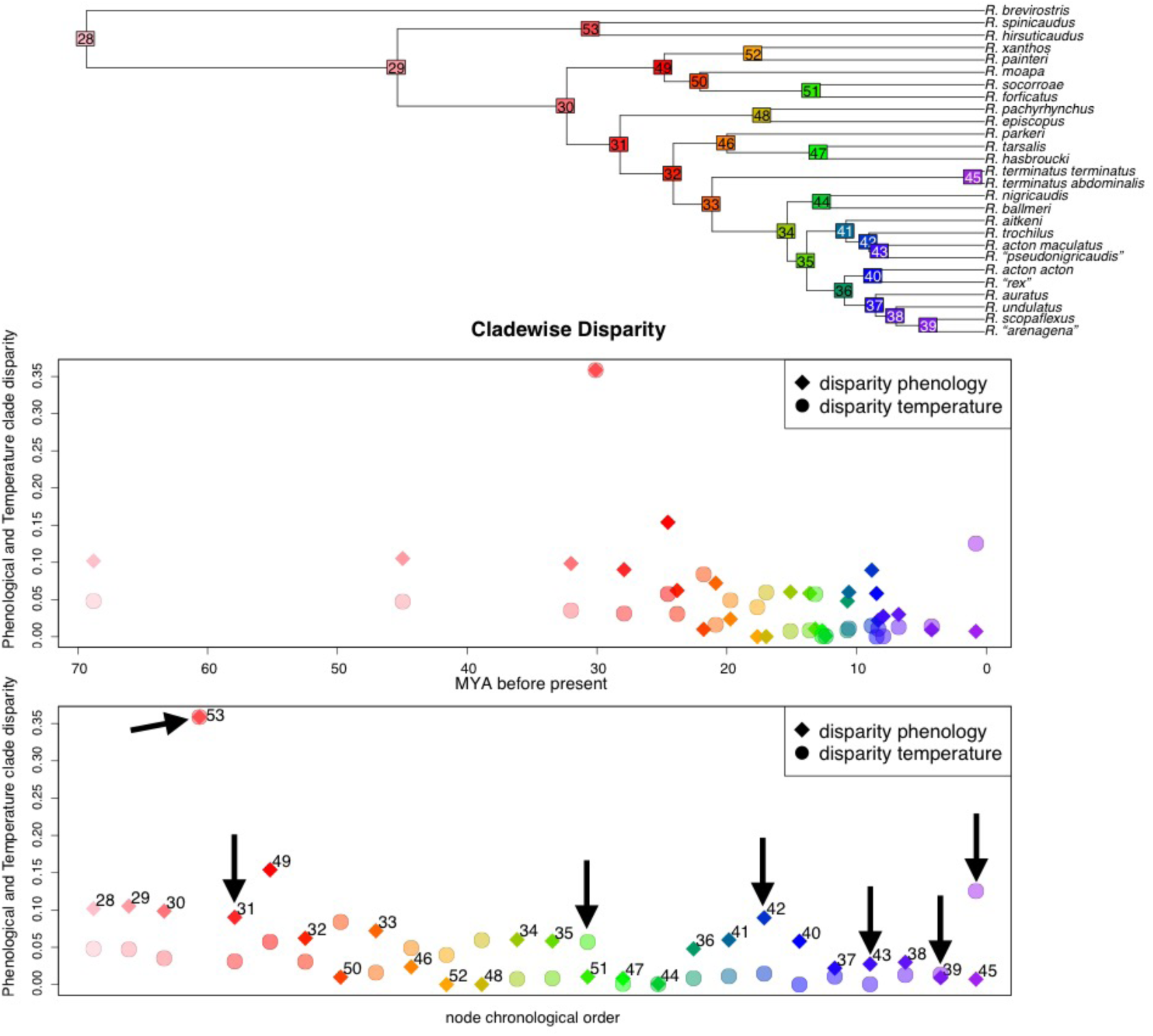
Cladewise disparity in phenology and temperature. **Top**, *Rhaphiomidas* chronogram with nodes colored by age, correlating to colors used in the panels below. **Middle**, temperature and phenological cladewise disparity of each subtree measured as average squared Euclidean distance. The Y-axis represents the cladewise disparity, and the X-axis is measured in millions of years. **Bottom**, temperature and phenological cladewise disparity of each subtree measured as average squared Euclidean distance. The Y-axis represents the cladewise disparity, and the X-axis is node chronological order, oldest to youngest. Arrows indicate nodes that are followed by a transition from light to dark coloration along one of their descendant branches.

**Figure 7.**
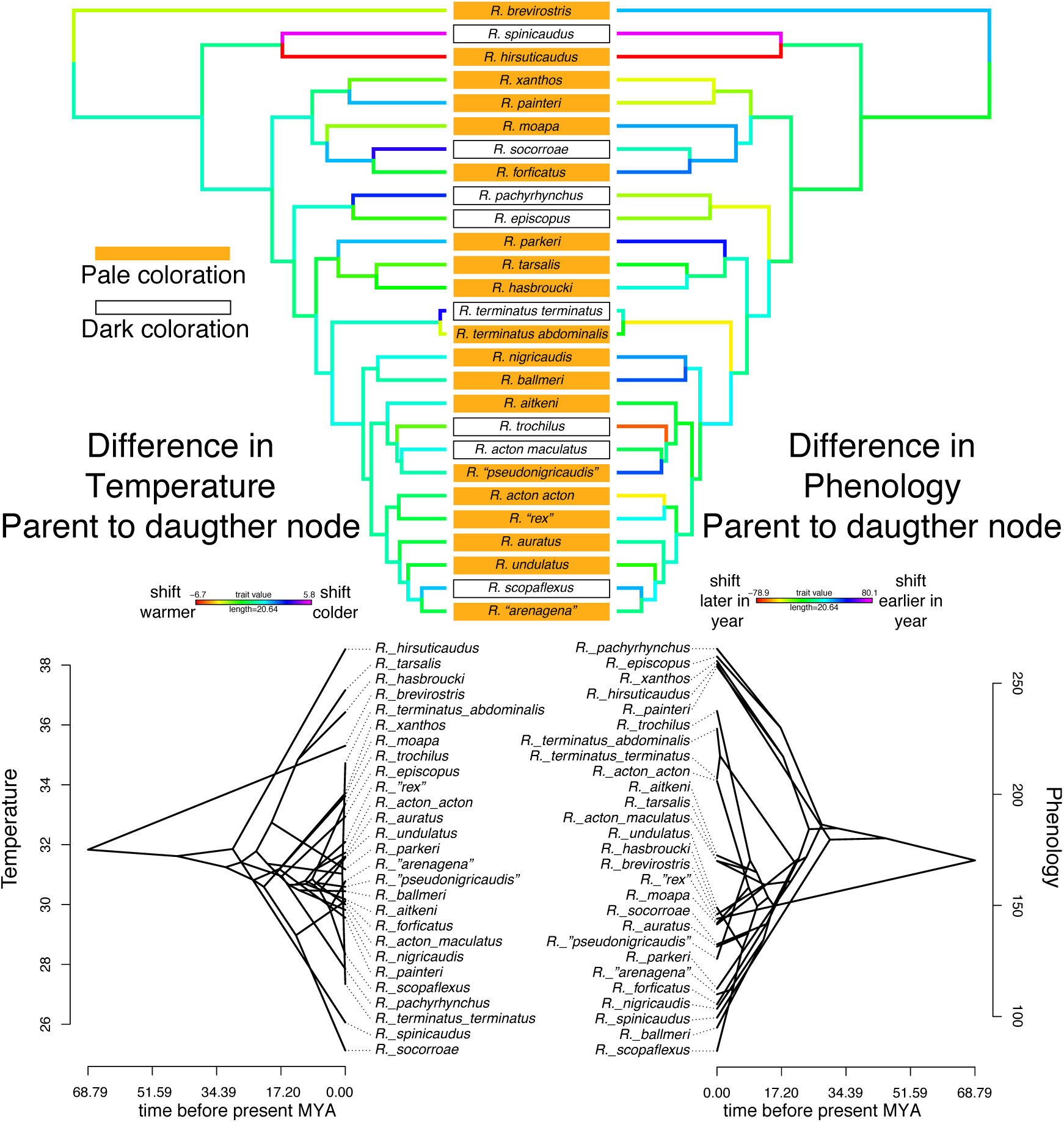
Color phenogram for mean daily maximum temperature (right half) and phenology (left half). Branch colors represent relative differences from parent to daughter nodes, similar to differences in the placement of parent to daughter nodes along the Y-axis as represented in the phenograms below, but phenogram nodes represent values not relative differences. Left half colors represent shifts to warmer or cooler temperatures between nodes (warmer colors = warmer shifts in temp., cooler colors = colder shifts in temp. relative to parent node), right half branch colors represent shifts in phenology (shifts later in year = warmer colors, earlier in year = cooler colors) relative to parent node. Orange rectangles indicate species with a light coloration, and black boxes indicate species with a dark coloration.

## DISCUSSION

Our results indicate that extracting climate data to match species’ phenological activity better exemplifies environmental conditions than the traditional Bioclim variables. Understanding the basic life history of an organism can go a long way toward producing more realistic niche models. Many animals, especially desert-adapted species, partition their life histories in response to local conditions and so share similar phenologies. For example, desert tortoises, *Gopherus agassizii,* spend the majority of their life underground in burrows, only active above ground for 153 hours per year (Nagy & Medica 1986). Modeling the effects of phenology on such species’ niches will be critical for better predictions of how these species might respond to future climate change. Additionally, our results show that *Rhaphiomidas* species adapt to different environments by evolving darker coloration (in colder conditions) and shifting phenologies.

### Using Accurate Temporal Data Improves Biological Insight

Our approach can be tailored to produce species specific environmental layers to best capture conditions that species are actually experiencing. This can be seen by examining the raw values obtained from the raster layers (Fig. 2). For example, when we look at a set of species that are partially sympatric but are separated temporally, the Bioclim data does not recover many differences between the species. By contrast, the Daymet data shows temperature values as completely separate or partially overlapping for both species. For example, *Rhaphiomidas sp. “arenagena”* is active earlier in the year compared to *R. sp. “rex”*. Their Bioclim BIO5 profiles overlap in temperature, whereas they do not overlap in the Daymet inference. This is seen with all the species that are partially sympatric but separated in time, as well as for some of the allopatric species that are also separated in time.

This result demonstrates a larger point that ecological inferences, such as testing for phylogenetic niche conservation of a trait related to temperature or another environmental variable, should not be made based on data that do not accurately link environmental data to the traits in question. By using environmental layers that are tailored to a species’ phenology in space and time, this will lessen misinterpretations of evolutionary trends, such as niche conservatism, due to the autocorrelation between geography and environment for species in allopatry (Warren et al. 2014). An abundance of data sources, such as Daymet, NDVI, and other MODIS data that are binned in daily or bi-weekly intervals, provide the much-needed data to make more accurate ecological and evolutionary inferences.

### Niche Shifts Through Color Evolution and Niche Tracking Through Changes in Phenology

*Rhaphiomidas* adapt to cooler conditions by evolving large areas of maculation (i.e. becoming darker), allowing them to emerge at times when conditions are less than optimal for them to fly efficiently or at all. Combining our *threshBayes* and *ancThresh* analyses, and the evidence from behavioral observations of activity (Cazier 1985; Kingsley 1996, 2002), we interpret the dark coloration of the cooler temperature species as an adaptation to warm themselves in order to take flight. This adaptation for dealing with cooler temperatures has allowed *Rhaphiomidas* to deviate from their thermal mean (as determined from ancestral state reconstructions) and allowed them to expand to new habitats.

Results from *threshBayes* analyses recover a significant and relatively strong correlation between temperature and coloration. This correlation is expected as *Rhaphiomidas* adults are usually only active above 26.6 ºC (Kingsley, 1996, 2002). Efficient warming for adequate flight is likely especially important for larger flies, such as *Rhaphiomidas* species, as their greater mass would require more time and energy to heat up. This phenomenon of darker coloration allowing ectotherms to warm themselves has been demonstrated in many taxa (Clusella Trullas et al., 2007; Svensson & Waller, 2013).

Our *ancThresh* results consistently recovered one ancestral node as dark in coloration – the *R. pachyrhynchus* and *R. episcopus* parent node. All other transitions in the tree were from pale to dark coloration, occurring only along the terminal branches, indicating that the observed dark coloration is not simply due to inheritance but instead a response to cooler environments. However, there are two dark *Rhaphiomidas* species that occur in relatively hot environments – *R. episcopus* and *R. trochilus*. In the case of *R. episcopus,* its dark coloration appears to be because the transition to this coloration occurred in the ancestor (Fig. 4). The reason the dark coloration is maintained in these species is not entirely clear and requires further investigation.

The *ancThresh* reconstructions in concert with the results from *contMap* can facilitate the identification of environmental niche tracking and/or adaptation to niches via phenology or coloration change. We found that all species were most likely pale in coloration ancestrally, indicating that changes are likely influenced by environment and not solely inheritance. We identified that some species evolved phenologies that are divergent relative to their sister taxon (Table 1). In addition, only one species experienced a shift in coloration solely as a response to cooler temperatures – *R. acton maculatus*. The other dark species changed both phenology and coloration, save *R. pachyrhynchus* and *R. episcopus,* which share a dark coloration ancestor. All other dark species, except *R. episcopus,* have denser setation (hairs), perhaps serving as insulation and providing additional morphological evidence of adaptation to a cool environment.

**Table 1.**
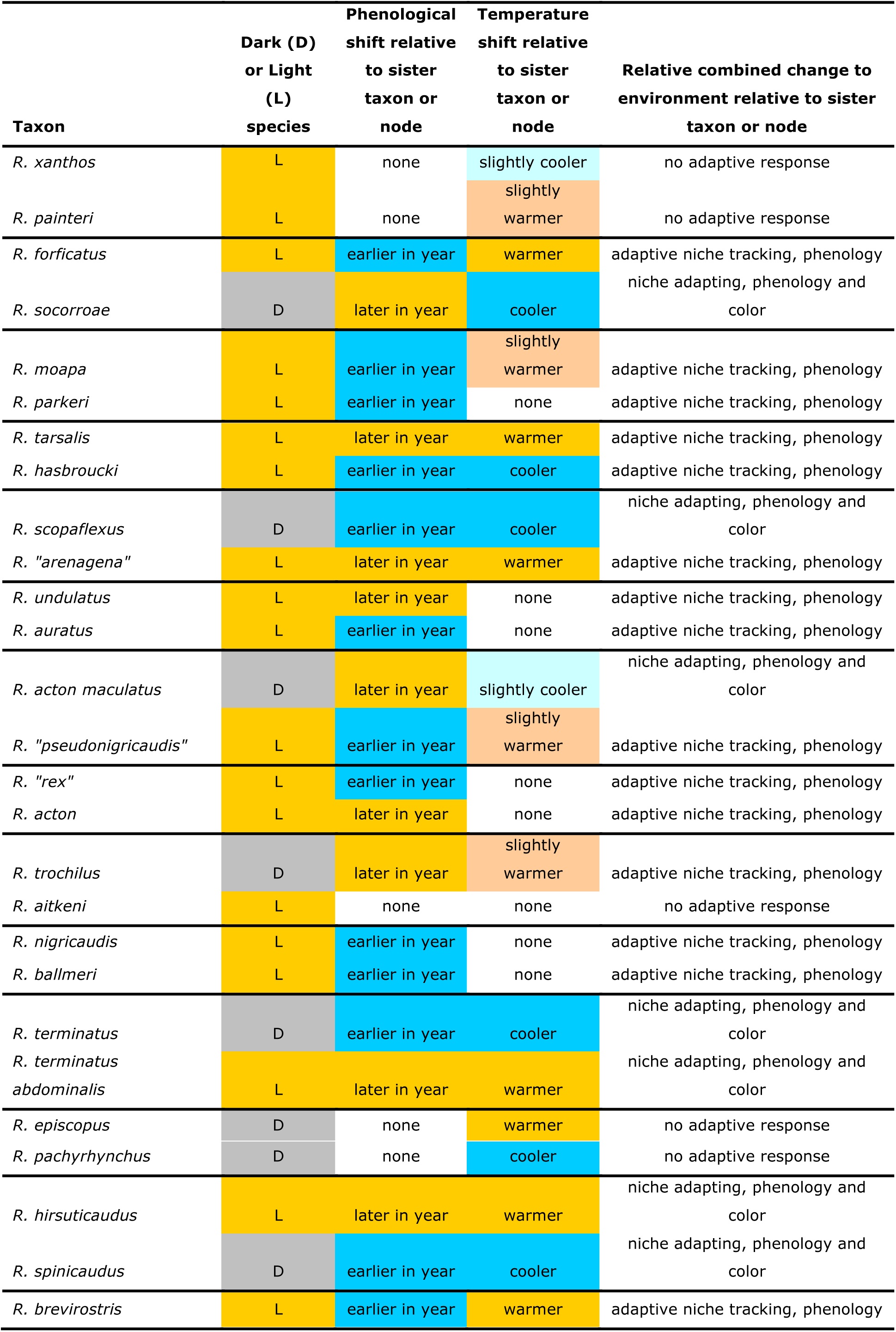
Summary of results identifying whether species adapt by tracking a niche by changing their phenology (adaptive niche tracking) or by strictly adapting to their niche through changing their coloration (niche adapting) or a combination of both. The factor(s) that evolved is/are listed after the adaptive response (niche tracking or niche adapting).

Our findings demonstrate that most *Rhaphiomidas* species tend to change their phenology as opposed to adapting their coloration to fit the climate, suggesting that phenology is a more labile trait. For example, *R. sp. “rex”* emerges about a month earlier than its sister taxon, *R. acton*. For both species, their phenologies coincide with late spring blooms in their different ranges (personal obs., M.H.V.D.). A phenology much later or earlier would render them unable to take advantage of these nectar resources. Their phenology should be a balance between resources availability and optimizing temperature. However, some species do not feed as adults, such as *R. hirsuticaudus* (Cazier 1985, personal obs. M.H.V.D.), so phenology should play an even more important role for such species to synchronize their emergence for reproduction.

## ACKNOWLEDGMENTS

The authors would like to thank Reviewer1, Reviewer2 and Sarah Crews for their careful comments that greatly improved the quality of the manuscript. The authors would also like to thank Jun Ying Lim for his help with statistical analyses and discussion of the paper. We also thank Greg Ballmer for his photos of *Rhaphiomidas* in Figure 1. We also thank Michelle Trautwein for her help in the writing of this manuscript and George Roderick, Rosemary Gillespie and Michael Balke for use of servers to perform the analyses. M.H.V.D. was funded by the following: Desert Research Fund, from The Community Foundation Serving Riverside and San Bernardino Counties, CA, USA; UC MEXUS Dissertation Research Grant; Theodore Roosevelt Memorial Fund from the American Museum of Natural History; Anza-Borrego Foundation and Institute (ABFI) Entomology; Committee for Research and Exploration of the National Geographic Society; Walker Fund in Entomology UCB; Robert L. Usinger Memorial Award UCB; NSF award DBI #1402102.

## SUPPORTING INFORMATION

Supporting Information link: https://figshare.com/s/0db26157b3b8c4cb3804, https://figshare.com/s/a680df82acaa692ecd79

## DATA ACCESSIBILITY

We have permanently uploaded all relevant data to figshare to make the research fully reproducible: Supporting Information link: https://figshare.com/s/0db26157b3b8c4cb3804 https://figshare.com/s/a680df82acaa692ecd79

## AUTHOR CONTRIBUTIONS

M.H.V.D. performed sampling, sequencing, phylogenetic dating, and wrote the first paper draft. A.J.R. wrote R scripts for Daymet data processing and PNO calculations, and M.H.V.D. and M.S.B. contributed to additional R scripts and informatics pipelines. All authors contributed to experimental design, analyses of the data, and final paper draft.

## BIOSKETCHES

**Matthew H. Van Dam** is interested in the interaction of geological processes and natural history of organisms in shaping biogeographical patterns; he is also interested in weevil systematics, including their evolution and improving phylogenomic methods.

**Andrew Rominger** is interested in how and why evolutionary history drives contemporary ecological dynamics. He approaches this challenge both from a theoretical perspective, building and testing synthetic theories of biodiversity based on statistical mechanics, and from an empirical perspective, collecting and analyzing new data about the evolution and ecology of arthropod diversity in island-like systems.

**Michael S. Brewer** uses genomics tools, bioinformatics, and arthropods as model organisms to answer broad evolutionary questions, especially those concerning the mechanisms that create and maintain biodiversity. Specific interests include taxonomy, biogeography, phylogenetics, and the evolution of complex traits (e.g., venomics, color evolution, and complex trait evolution).

